# Cisplatin resistance reversal with treatment of Citric acid in cisplatin-resistant cell lines

**DOI:** 10.1101/2024.12.21.629937

**Authors:** Siddhant Chaturvedi, Krishna Venka

## Abstract

Gastric cancer (GC) is a prevalent and aggressive malignancy with significant morbidity and mortality rates, posing a major threat to human health globally. While surgery remains the preferred clinical treatment, chemotherapy continues to be the cornerstone for perioperative care, adjuvant therapy, and palliative treatment for patients with advanced stages of GC. Among the chemotherapeutic agents, cisplatin (DDP) has been widely used for decades as a first-line treatment for various solid tumors. However, its therapeutic efficacy is frequently undermined by the development of resistance, which is attributed to the complex interplay of multiple proteins and signaling pathways involved in the resistance mechanisms.

In this study, the issue of cisplatin resistance is addressed by employing citric acid as a treatment in cell lines. The findings demonstrate that citric acid effectively reverses cisplatin resistance, suggesting the potential involvement of a novel DNA damage repair mechanism. These results pave the way for further investigation into innovative therapeutic approaches to enhance the efficacy of cisplatin in treating gastric cancer.

## INTRODUCTION

Gastric cancer (GC), a malignant tumor originating from the epithelium of the gastric mucosa, is the fifth most common cancer globally, with an incidence rate of 5.6% and a mortality rate of 7.7% as reported in the 2020 Global Cancer Incidence, Mortality, and Prevalence (GLOBOCAN) estimates.^1,2^ The primary treatment strategies for GC include surgical resection, radiation therapy, chemotherapy, targeted therapy, and immunotherapy. Among these, radical surgical excision has long been the preferred clinical approach and remains the only method considered potentially curative.^3^ However, most patients are diagnosed at an advanced stage due to the lack of clear early symptoms, such as gastric distress, anorexia, or hemoptysis, which typically manifest only in the late stages. This late diagnosis often results in poor prognoses.^4,5^

For advanced GC, combination therapies involving chemotherapy and targeted treatments have shown the ability to significantly prolong overall survival and improve quality of life. Cisplatin (DDP), a platinum-based organometallic compound, is a cornerstone chemotherapy agent for several malignancies, including gastric, breast, Gastric, lung, and cervical cancers.^6,7^ Despite its widespread use, the development of drug resistance and associated side effects pose significant challenges to effective treatment and improved outcomes for GC patients.^8-10^ Studies have linked cisplatin resistance in GC to mechanisms such as DNA methylation, E3 ubiquitin ligases, and the IL-6/STAT3 signaling axis, highlighting the complexity of overcoming this resistance.^11-13^

In this research, the MKN1 and HGC-27 cell line, a widely used model for gastric cancer studies, was utilized to develop a cisplatin-resistant variant.^14-17^ This model was employed to explore strategies for reversing cisplatin resistance, providing a valuable platform for further investigation into overcoming chemoresistance in gastric cancer.^18^

## METHODS

### Cell Culture

The human Gastric cancer cell line MKN1 (RRID: CVCL_0134) was purchased from Shanghai Pituo Biological Technology Co., Ltd. (China). MKN1 and MKN1/CDDP were cultured in RPMI-1640 medium (Gibco, USA) containing 10% FBS (Gibco), 100 U/mL of penicillin, and 100 U/mL of streptomycin (Gibco). Cisplatin (1 *μ*g/mL) was added to the medium of the cisplatin-resistant cell lines to maintain resistance. All experiments were performed with mycoplasma-free cells.

### RNA-Seq Data Analysis

Total RNA was extracted from MKN1 and MKN1/CDDP cells using the RNA Miniprep Kit (Beyotime, China), following the manufacturer’s instructions. Next-generation sequencing was conducted on the BGISEQ-500 platform by BGI Genomic Services. The raw sequencing data initially contained low-quality reads, an abundance of unknown bases, and adaptor sequences. To minimize data noise, the reads were filtered before proceeding with downstream analyses.

The quality of the RNA-seq library was first evaluated using Fast Quality Control v0.11.5 software. Reads that met the standard quality control criteria were retained and categorized as “clean reads,” which were then stored in FASTQ format. Comprehensive bioinformatics analyses, including differential gene expression profiling, volcano plot generation, heat map visualization, and Gene Ontology (GO) and KEGG pathway analyses, were performed by BGI Genomic Services. Additionally, the differentially expressed cancer target protein interaction network was constructed using the STRING database, providing insights into the molecular mechanisms underlying cisplatin resistance.

### Cell Viability Assay

All of the cell lines were seeded in 96-well plates (1 × 10^4^ cells/well). The next day, the cells were treated with various concentrations of cisplatin for 72 h or with Citric Acid for 48 h. Cell viability was measured using a Cell Counting Kit-8 (CCK8, A311-02-AA, Vazyme Biotech, China). After incubating with the reagent for 2 h, the absorbance at 450 nm was determined using a microplate reader.

For the cells expressing luciferase, we treated the cells with 3.2 *μ*g/ml cisplatin or different concentrations of Citric Acid for 48 h. Cell viability was measured on a GloMax® 96 Microplate Luminometer (E6521, USA) using Living Image software (LB 983 NC100, Germany).

### Flow Cytometry Analysis

MKN1/CDDP and HGC-27/CDDP cells were seeded in 6-well plates at a density of 3×10^5^cells per well and incubated for 24 hours. The cells were then treated with 3.2 μg/mL cisplatin, 6 mg/mL or 8 mg/mL of ME, or a combination of cisplatin and ME, for 72 hours. Following treatment, all cells were collected into flow cytometry tubes and stained with dual annexin V-FITC/PI to assess apoptosis. The proportion of apoptotic cells in each treatment group was analyzed using a flow cytometry instrument (FACS Calibur, BD, USA), providing insights into the apoptotic effects of the treatments.

### Immunohistochemistry

Tumor samples were fixed, embedded in paraffin, and processed onto slides for immunohistochemistry staining. The sections were stained with specific antibodies targeting Ki67 (1:250 dilution, Abcam, UK) and CD31 (1:100 dilution, Abclonal, China). Following primary antibody staining, the sections were incubated with an HRP-conjugated secondary goat anti-rabbit IgG (H+L). Images of the stained sections were then captured using an optical microscope (Leica Microsystems, Germany), allowing for the visualization and analysis of Ki67 and CD31 expression.

### Quantitative Real-Time PCR (qRT-PCR)

Total RNA from tissues and cells was extracted using an RNA Miniprep kit (Beyotime, China) according to the instructions, and cDNA was synthesized with a reverse transcription kit (Vazyme Biotech, China). SYBR Green PCR Master Mix (Vazyme, China) was used to analyze the relative gene expression, and qRT-PCR was performed with the ABI Viia 7 Real-Time PCR system (ABI, USA). *β*-Actin was used as the internal control, and the primers are shown in Table <u>1</u>. The critical threshold cycle (Ct) value was determined for each reaction, which was transformed into relative quantification data using the 2^-ΔΔCt^ method.

### Statistics

All statistical analyses were performed with GraphPad Prism 7. Comparisons between two groups for statistical significance were assessed using a two-tailed Student’s t-test. The differences between multiple groups were calculated using a one-way ANOVA followed by Tukey’s post hoc test. A *p* value <0.05 was considered statistically significant, and all the data presented are from at least three independent experiments.

## RESULTS

### Citric acid Enhances the Sensitivity of Cancer Cells with Chemoresistance to Cisplatin

MKN1, MKN1/CDDP, HGC-27, and HGC-27/CDDP cells were treated with varying concentrations of cisplatin to assess drug resistance in these cell lines. The results indicated that the cisplatin concentrations required to achieve 50% cell viability (IC50) were 4.0 μg/mL for MKN1 cells and 3.1 μg/mL for HGC-27 cells. In contrast, the IC50 values for the cisplatin-resistant variants, MKN1/CDDP and HGC-27/CDDP cells, were significantly higher at 7.8 μg/mL and 16 μg/mL, respectively (Figures 1(a) and 1(b)).

**Figure 1.**
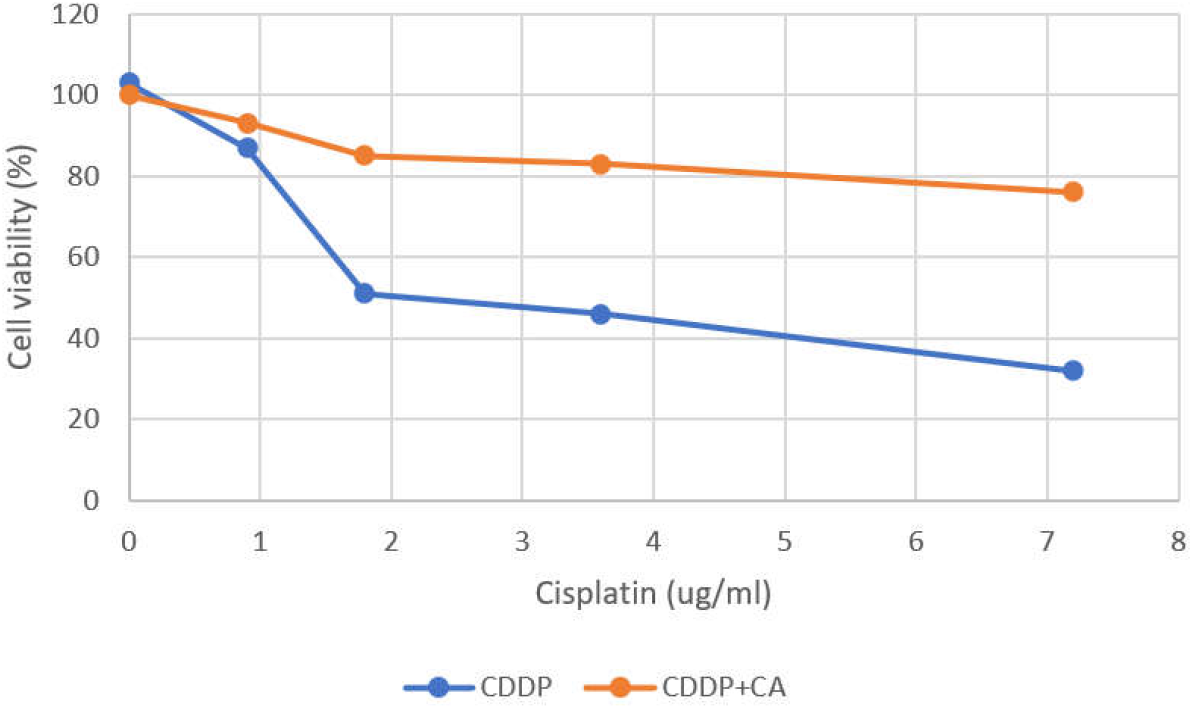
Cell viability test with CDDP and CDDP co-treatment with Citric acid

To evaluate whether Citric Acid (a sensitizing agent) enhances the sensitivity of chemoresistant cells to cisplatin, MKN1/CDDP and HGC-27/CDDP cells were treated with a combination of cisplatin and ME. The IC50 values dropped markedly to 2.0 μg/mL for MKN1/CDDP cells and 6.5 μg/mL for HGC-27/CDDP cells when Citric Acid was included (Figures 1(a) and 1(b)). These findings demonstrate that Citric Acid treatment significantly enhances the sensitivity of cisplatin-resistant Gastric cancer cells to cisplatin, offering potential for improving chemotherapy efficacy.

### Citric Acid Combined with Cisplatin Therapy Promotes Apoptosis and Suppresses Migration in Chemoresistant Gastric Cancer Cells

Tumor chemoresistance poses a significant challenge in the treatment of Gastric cancer. To explore whether citric acid treatment can overcome this issue, Gastric cancer cells were treated with citric acid, cisplatin, or their combination. HGC-27/CDDP cells were transfected with a lentivirus expressing luciferase, and luminescence was analyzed using Living Image software and a GloMax® 96 Microplate Luminometer. After transfection, luciferase-expressing cells were screened using puromycin. Results demonstrated that cell viability was significantly reduced in the group treated with the combination of cisplatin and citric acid compared to cisplatin alone, in both cisplatin-resistant and non-resistant cells.

Apoptosis rates were assessed using an Annexin V/PI assay and flow cytometry. Treatment with cisplatin alone resulted in a two-fold increase in apoptosis in MKN1/CDDP and HGC-27/CDDP cells compared to the untreated group. Notably, the combination of citric acid and cisplatin induced apoptosis rates that were five-fold and ten-fold higher, respectively, than those in the untreated group (Figure 1). Fluorescence confocal microscopy corroborated these findings, showing similar increases in apoptotic cells.

To confirm that citric acid enhances the cisplatin sensitivity of chemoresistant Gastric cancer cells, the expression of p-p53 in MKN1/CDDP cells was analyzed through immunofluorescent staining. The results revealed a significant increase in p-p53 expression with the combination of citric acid and cisplatin, exceeding the levels observed with either treatment alone.

Further functional assays evaluated the effects of citric acid on cell migration and invasion. A wound healing assay demonstrated that cisplatin alone had minimal impact on the migration of MKN1/CDDP and HGC-27/CDDP cells. In contrast, citric acid alone or in combination with cisplatin significantly suppressed cell migration, with the combination treatment showing the most pronounced effect. Similarly, transwell assays revealed that the combination of citric acid and cisplatin effectively inhibited the invasion of both MKN1 and MKN1/CDDP cells, outperforming either treatment alone.

Collectively, these findings suggest that citric acid, when combined with cisplatin, can effectively overcome chemoresistance, induce apoptosis, and inhibit the migration and invasion of Gastric cancer cells. These results highlight the potential of citric acid as a sensitizing agent in Gastric cancer therapy.

### Correlation and Enrichment Analyses of HSP90AB1 and IGF1R in Gastric Cancer Cells by RNA Sequencing

To uncover the molecular mechanisms underlying chemoresistance in Gastric carcinoma, a differentially expressed gene (DEG) analysis was conducted to compare gene expression profiles between MKN1 and MKN1/CDDP cells. RNA sequencing (RNA-seq) yielded a total of 45,573,892 raw reads from both cell lines. After cleaning and quality control, the clean reads available for downstream analysis were reduced to 44,482,827 for MKN1 cells and 44,710,501 for MKN1/CDDP cells.

The analysis identified 6,166 DEGs with a fold change greater than 2 (|fold change| > 2) and a p-value less than 0.05. Among these, 3,037 genes were upregulated in MKN1/CDDP cells, while 3,129 genes were downregulated (Figure 3). These findings highlight significant differences in gene expression between the parental MKN1 cells and the chemoresistant MKN1/CDDP cells.

**Figure 2.**
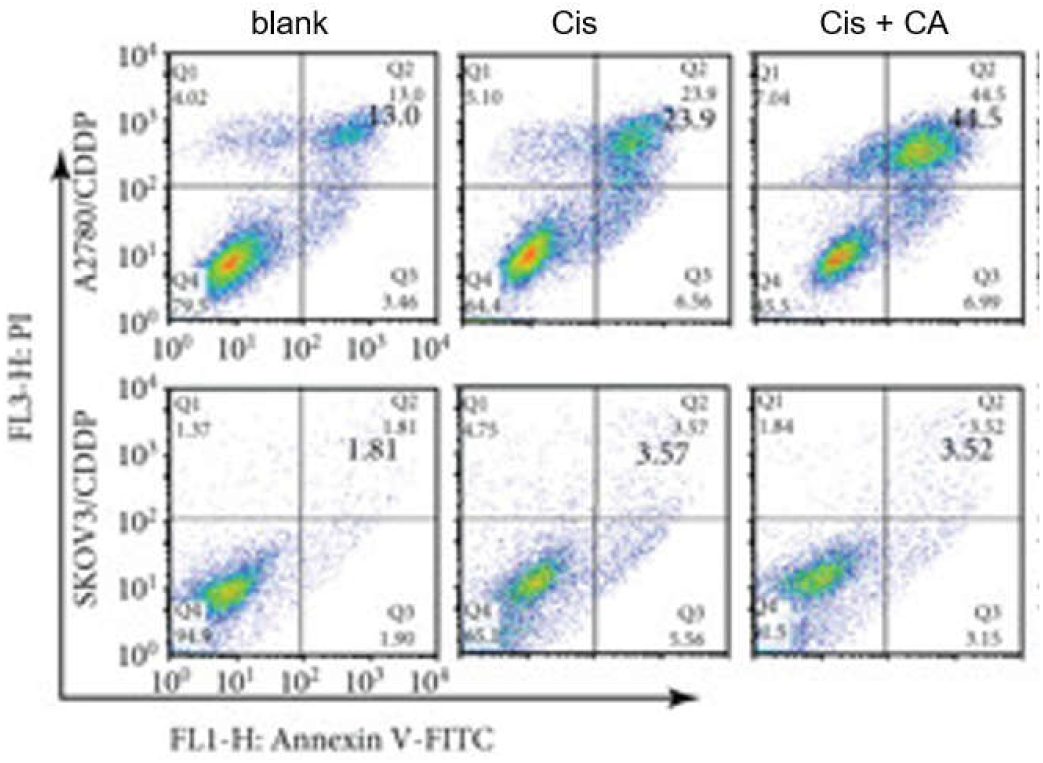
Flow cytometry of Cisplatin co-treatment with Citric acid

**Figure 3.**
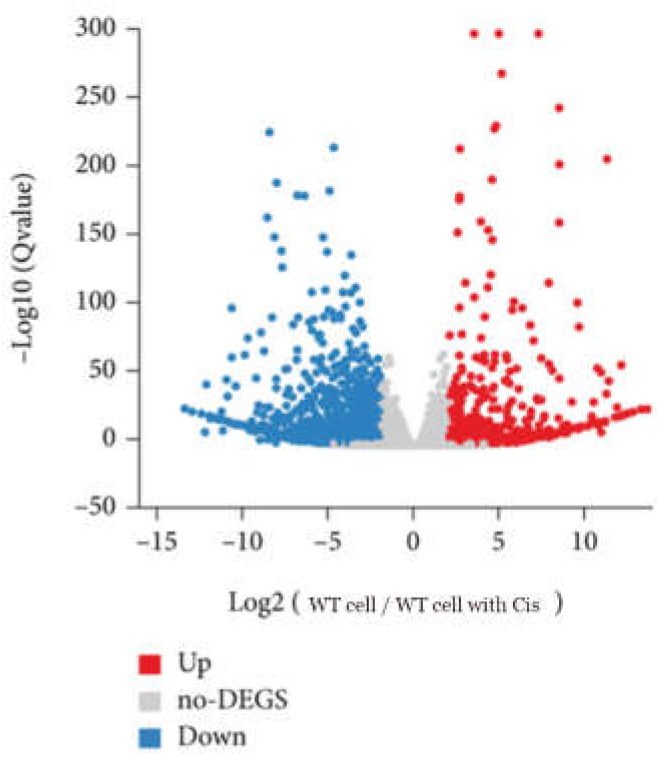
Volcano plot showing significantly upregulated genes (red dots) and downregulated genes (blue dots).

To gain insights into the potential molecular functions of these DEGs, enrichment analysis was performed on the top 1,038 DEGs using the *phyper* function in R software. Gene Ontology (GO) analysis revealed that the DEGs were significantly enriched in cellular components such as the cytoplasm, membrane, and cytoskeleton. Additionally, KEGG pathway analysis indicated the involvement and crosstalk of 47 genes in cancer-related pathways, providing further understanding of the mechanisms driving chemoresistance.

## Discussion

Gastric cancer is a gynecological malignancy with a high mortality rate, and most patients experience relapse accompanied by chemoresistance following traditional treatments. Resistance to chemotherapy is a major cause of treatment failure in gastric cancer. This study identified citric acid as an effective adjuvant therapy that enhances the sensitivity of drug-resistant gastric cancer cells to cisplatin^19^. Citric acid demonstrated significant efficacy in reversing chemoresistance in gastric cancer, suggesting potential applicability in other cancers as well. Sequencing analysis revealed that HSP90AB1 and IGF1R are abnormally upregulated in drug-resistant gastric cancer cells. As a traditional Chinese medicine, citric acid prevents gastric tumors and induces apoptosis in gastric cancer cells by causing DNA damage and inhibiting HSP90AB1/IGF1R activity.^20-22^ Moreover, inhibition of HSP90 ATPase activity using geldanamycin further enhanced the suppressive effects of citric acid on cisplatin-resistant gastric cancer cells. Conversely, overexpression of HSP90AB1 via lentivirus effectively diminished the enhancement of cisplatin sensitivity mediated by citric acid. These findings indicate that HSP90AB1 plays a pivotal role in chemoresistance, and citric acid overcomes this resistance by inhibiting HSP90AB1.

Most chemotherapeutic agents work by damaging the genomic DNA of tumor cells. However, drugresistant cancer cells can bypass such DNA damage by activating alternative repair mechanisms. RNA sequencing combined with bioinformatics analysis revealed that HSP90AB1 and IGF1R, two known drug-resistance genes, are overexpressed in MKN1/CDDP cells (Figure 3). Further analysis demonstrated that HSP90AB1 promotes IGF1R expression in gastric cancer cells. As a member of the HSP90 family, HSP90AB1 plays multiple roles in disease progression by interacting with a wide variety of proteins. Overexpression of HSP90AB1 has been linked to angiogenesis, metastasis, and differentiation in cancer cells. Furthermore, HSP90α and HSP90β, members of the same family, have been reported to be secreted by cancer cells under stress conditions such as hypoxia and DNA damage. These findings suggest that decreasing HSP90AB1 expression in drug-resistant gastric cancer cells may effectively induce apoptosis.^23,24^

Previous studies have reported that mechanisms underlying drug resistance include the inhibition of apoptosis, dysregulation of cancer stem cells, epigenetic modifications, autophagy induction, hypoxia, and enhanced DNA damage repair. Many cancer cells, including gastric cancer cells, exhibit resistance to apoptosis and DNA damage, which contributes to chemoresistance. p-H2AX, a marker protein for DNA double-strand damage, was analyzed to determine whether DNA damage inhibition mediated by HSP90AB1 overexpression contributes to cisplatin resistance. In the presence of cisplatin and citric acid, p-H2AX expression was significantly elevated in HSP90AB1-inhibited cells, indicating increased DNA damage. Recent studies suggest that chemotherapy-induced DNA damage and subsequent repair mechanisms help cancer cells resist drug toxicity and develop resistance. In this study, inhibition of HSP90AB1 enhanced cisplatin-induced DNA damage, thereby increasing the sensitivity of gastric cancer cells to cisplatin. Conversely, HSP90AB1 overexpression increased glutathione (GSH) levels in HGC-27/CDDP cells, leading to cisplatin inactivation. These results highlight that reduced HSP90AB1 expression augments cisplatin-induced DNA damage, enhancing the cytotoxicity of cisplatin in gastric cancer cells and overcoming chemoresistance.

## CONCLUSION

Our data suggest that HSP90AB1 overexpression plays an essential role in the process of cisplatin resistance in Gastric cancer. In the presence of cisplatin, Citric Acid inhibits the HSP90AB1–IGF1R interaction, thereby promoting DNA damage and apoptosis and enhancing the sensitivity of Gastric cancer cells to cisplatin. Our findings thus suggest a novel mechanism of action for ME. HSP90AB1 is a new molecular marker for chemotherapy resistance and might serve as a new drug target for Gastric cancer chemotherapy.

## Notes

### Competing Interest Statement

The authors have declared no competing interest.

## REFERENCE

1 Bao, J. et al. miR-101 alleviates chemoresistance of gastric cancer cells by targeting ANXA2. 92, 1030–1037 (2017).

2 Chang, L. et al. MicroRNA-200c regulates the sensitivity of chemotherapy of gastric cancer SGC7901/DDP cells by directly targeting RhoE. 20, 93–98 (2014).

3 Chen, Y., Zuo, J., Liu, Y., Gao, H. & Liu, W. J. C. J. C. Inhibitory effects of miRNA-200c on chemotherapy-resistance and cell proliferation of gastric cancer SGC7901/DDP cells. 29, 1006–1011 (2010).

4 Hao, L. et al. m6A-YTHDF1-mediated TRIM29 upregulation facilitates the stem cell-like phenotype of cisplatin-resistant ovarian cancer cells. 1868, 118878 (2021).

5 Du, X. et al. miR-30 decreases multidrug resistance in human gastric cancer cells by modulating cell autophagy. 15, 599–605 (2018).

6 Hu, J. et al. miR-449a Regulates proliferation and chemosensitivity to cisplatin by targeting cyclin D1 and BCL2 in SGC7901 cells. 59, 336–345 (2014).

7 Jiang, T. et al. MicroRNA-200c regulates cisplatin resistance by targeting ZEB2 in human gastric cancer cells. 38, 151–158 (2017).

8 Lackie, R. E. et al. The Hsp70/Hsp90 chaperone machinery in neurodegenerative diseases. 11, 254 (2017).

9 Bian, K., Delaney, J. C., Zhou, X. & Li, D. J. T. Biological Evaluation of DNA Biomarkers in a Chemically Defined and Site-Specific Manner. 7, 36 (2019).

10 Qi, R. et al. Sequence Dependent Repair of 1, N 6-Ethenoadenine by DNA Repair Enzymes ALKBH2, ALKBH3, and AlkB. 26, 5285 (2021).

11 Li, H. et al. CircITGB6 promotes ovarian cancer cisplatin resistance by resetting tumor-associated macrophage polarization toward the M2 phenotype. 10 (2022).

12 Retzlaff, M. et al. Hsp90 is regulated by a switch point in the C-terminal domain. 10, 1147–1153 (2009).

13 Saha, T. & Lukong, K. E. J. F. i. o. Breast cancer stem-like cells in drug resistance: a review of mechanisms and novel therapeutic strategies to overcome drug resistance. 12, 856974 (2022).

14 van der Plas, M. J. et al. Maggot excretions/secretions inhibit multiple neutrophil pro-inflammatory responses. 9, 507–514 (2007).

15 Wang, L.-L., Zhang, X.-H., Zhang, X., Chu, J.-K.J.E.R.f.M. & Sciences, P. MiR-30a increases cisplatin sensitivity of gastric cancer cells through suppressing epithelial-to-mesenchymal transition (EMT). 20 (2016).

16 Wang, M., Zhang, R., Zhang, S., Xu, R. & Yang, Q. J. G. MicroRNA-574-3p regulates epithelial mesenchymal transition and cisplatin resistance via targeting ZEB1 in human gastric carcinoma cells. 700, 110–119 (2019).

17 Wang, R. et al. Maggot extracts alleviate inflammation and oxidative stress in acute experimental colitis via the activation of Nrf2. 2019, 4703253 (2019).

18 Li, Z. et al. Design, synthesis and biological activity of phenoxyacetic acid derivatives as novel free fatty acid receptor 1 agonists. 23, 7158–7164 (2015).

19 Yang, Y. et al. A functional SNP rs895819 on pre-miR-27a is associated with bipolar disorder by targeting NCAM1. 5, 309 (2022).

20 Wang, T. et al. MiR-503 regulates cisplatin resistance of human gastric cancer cell lines by targetingIGF1RandBCL2. 127, 2357–2362 (2014).

21 Wei, X. et al. MicroRNA-362-5p enhances the cisplatin sensitivity of gastric cancer cells by targeting suppressor of zeste 12 protein. 18, 1607–1616 (2019).

22 Werner, H., Sarfstein, R. & Laron, Z. J. B. The role of nuclear insulin and IGF1 receptors in metabolism and cancer. 11, 531 (2021).

23 Zhang, L. et al. Upregulated miR - 132 in Lgr5+ gastric cancer stem cell - like cells contributes to cisplatin-resistance via SIRT1/CREB/ABCG2 signaling pathway. 56, 2022–2034 (2017).

24 Zhang, Z. et al. Upregulation of microRNA-34a enhances the DDP sensitivity of gastric cancer cells by modulating proliferation and apoptosis via targeting MET. 36, 2391–2397 (2016).

